# A Generative Growth Model for Thalamocortical Axonal Branching in Primary Visual Cortex

**DOI:** 10.1101/288522

**Authors:** Pegah Kassraian Fard, Michael Pfeiffer, Roman Bauer

## Abstract

Axonal morphology displays large variability and complexity, yet the canonical regularities of the cortex suggest that such wiring is based on the repeated initiation of a small set of genetically encoded rules. Extracting underlying developmental principles can hence shed light on what genetically encoded instructions must be available during cortical development. Within a generative model, we investigate growth rules for axonal branching patterns in cat area 17, originating from the lateral geniculate nucleus of the thalamus. This target area of synaptic connections is characterized by extensive ramifications and a high bouton density, characteristics thought to preserve the spatial resolution of receptive fields and to enable connections for the ocular dominance columns. We compare individual and global statistics, such as a newly introduced asymmetry index and the global segment-length distribution, of generated and real branching patterns as the benchmark for growth rules. We show that the proposed model surpasses the statistical accuracy of the Galton-Watson model, which is the most commonly employed model for biological growth processes. In contrast to the Galton-Watson model, our model can recreate the log-normal segment-length distribution of the experimental dataset and is considerably more accurate in recreating individual axonal morphologies. To provide a biophysical interpretation for statistical quantifications of the axonal branching patterns, the generative model is ported into the physically accurate simulation framework of Cx3D. In this simulation environment we demonstrate how the proposed growth process can be formulated as an interactive process between genetic growth rules and chemical cues in the local environment.

## 1 Introduction

The canonical regularities of cortex imply that axonal branching patterns, although displaying complex and highly variable morphologies, have developed based on the repeated initiation of a small set of genetically encoded rules. Extracting underlying developmental principles can hence shed light on what genetically encoded instructions must be available during cortical development [26]. In particular, we assess here developmental rules for axonal arborizations in cat area 17, originating from laminae A of dorsal lateral geniculate nucleus (LGN) of the thalamus. Axonal ramifications in this region are characterized by a high bouton density and extensive arborizations [75], and are here due to their morphology referred to as *florets.* These arborizations must be organized with high precision to preserve the spatial resolution of the associated receptive fields [2], and to enable dense synaptic connections for the ocular dominance columns [32, 33]. Nevertheless, the morphology of these arborizations is highly variable, reaching from florets consisting of only a single branch to elaborate structures with tens of segments of highly diverse lengths. The motif of canonical regularities observed in the cortex however suggests that also the development of the florets is based on a small set of such rules, giving rise to these elaborate arborizations by their repeated initiation in interaction with the local environment [4–8,17,42,48]. As neural computations - opposed to computation in artificial systems - arise within a self-organized system, understanding the principles guiding cortical development promises an understanding of the information processing the brain is capable of.

Models of local axonal growth have been few in number, and so far based on simple Galton-Watson branching processes [7, 45]. The Galton-Watson model [29], the best established and most studied growth model, allows at each step for either bifurcation, growth or halting, all with constant probabilities during the entire growth process. It does however not incorporate biophysical constraints, and generates an exponential distribution of segment-lengths, failing to match a crucial metric of neurites with other segment-length distributions. The florets in cat area 17 for instance possess a log-normal segment-length distribution, a length-distribution characteristic for a variety of growth processes [1, 7].

Other models of neurite growth are mostly dedicated to dendritic outgrowth [13, 62, 68, 71]. Classical dendrite growth models typically assume different growth rules depending on the growth cones’ position in the topology of the neurite [62, 68, 71], an assumption opposed to biological principles of local autonomy [5, 27, 73].

Another approach to the modelling of neurite morphologies has been inspired by Cajal’s conversation principles of cytoplasmic volume, space and conduction time [9, 10, 12, 13]. Realistic recreations of diverse types of dendrite morphologies have been achieved for instance by combining constraints on the total wiring length with constraints on the path length to the root of the dendrite [13]. Besides being mostly concerned with dendrite morphologies [13], this formal optimization approach does not allow for insights into the mechanistic underpinnings of the neurite growth process. The global optimization requirement stands furthermore in contrast to biological principles of local autonomy, and does not take environmental cues into account.

We propose here a generative axonal growth model which is based on principles of cortical development and provides a mechanistic account of the floretal growth process. We assess the ability of the proposed model to reproduce individual and global statistics of the dataset as a benchmark for the suggested growth rules. The model is optimized for the global segment-length distribution and a weighted morphological asymmetry quantification. The later extends the classically used asymmetry index [11, 67] to include metric properties, allowing for a finer control of morphological asymmetries of generated florets. In a further step, we port the generative model into the simulation environment of cortical development *Cx3D*, which allows us to provide a biophysical interpretation of the observed statistics. Here we model the morphological asymmetry of the florets as the result of interactions of the growth process with a chemotaxic cue, crucial for axonal path finding [19, 22, 41], in the local environment. This translation underlines the relevance of the suggested growth rules in light of neural computation and allows for integration of the proposed neurite wiring model as part of broader neocortical genesis [73]. We show here that our generative model, which makes use of only a small set of simple rules, yields well-matched individual and global statistics of thalamocortical axonal branching in the target area.

## 2 Materials and Methods

### 2.1 Dataset

The neurons examined in this study were obtained from anesthetized adult cats that had been prepared for in vivo intracellular recording. All experiments were carried out by Kevan A.C. Martin and colleagues under the authorization of animal research licenses granted by the Home Office of the U.K. and the Cantonal Veterinary Authority of Zurich. A total of 426 florets from 10 axons was collected from 5 adult cats. All axons originate in A laminae of the dorsal part of the lateral geniculate nucleus and project to layer 4 of area 17. Axons were classified as X or Y-type using a battery of tests [23, 35]. Three of these axons have been used in previous studies [2,6–8]. Surgical details are found in [14]. After labelling the axons in the anaesthetized cat in vivo using intracellular injections of horseradish peroxidase (HRP) or anterograde tracer biotinylated dextran amine (BDA), the axons were reconstructed from blocks of histologically prepared and serially sectioned tissue. Using a x40 or x100 oil immersion objective attached to a light microscope and drawing tube the axons were reconstructed in 2D and 3D by using either the in-house 3-D reconstruction system TRAKA or Neurolucida. The reconstructions were stored as a list of data points for further usage. Florets from a single axon branching in layer 4 of the visual cortex are shown in Figure 1 panel B. Figure 1 panel C shows various florets aligned by their root segment. We define the *segment length* as the length of the neurite between branching points, or between branching points and the tip of a floret, measured in microns [μm] in 3D space. Among the 426 reconstructed florets, 260 are *trivial florets* which consist of a single segment.

**Figure 1:**
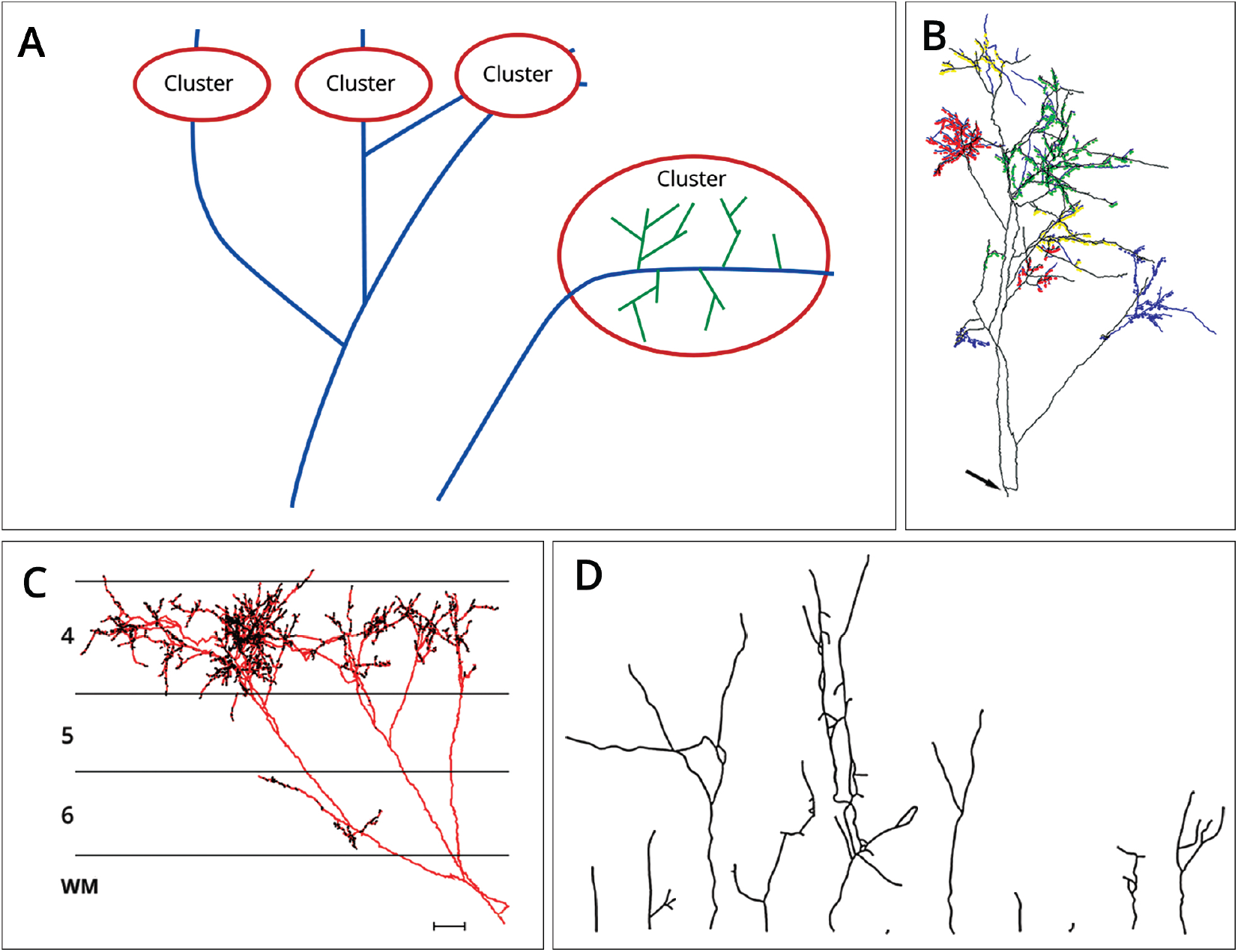
(A) Clusters containing florets (green) and long-range axonal projections (blue). Besides a higher branching probability than present at the axonal trunk, florets can be characterized by a higher bouton density. The long-range axonal parts outside of the clusters in contrast display fewer branching occurrences, longer branches, and a lower bouton density [8, 75]. (B) Labeled Y axon terminals of cat primary visual cortex arborising in layers 4 and 6. The axon was reconstructed from serial sections and has been rotated to optimize the view of the coloured florets. The axon emerges from the white matter at the location indicated by the arrow. (C) Displayed is the same axon (red) as in (B), here however from a coronal view. The axon emerges from white matter, here indicated by ‘WM’. The terminals of the axon focussing on layers 4 and 6 are indicated in black. A horizontal 100 micron scale bar appears in the bottom right hand corner. (D) Florets aligned by their root segments. Floret morphology is highly variable, with florets consisting of only a single segment (‘trivial florets’) as well as larger florets with many and highly diverse segments being present in the dataset. All experimental data is courtesy of Kevan A.C. Martin and colleagues, Institute of Neuroinformatics, Zurich.

### 2.2 Definition of Florets

Extensive ramifications in the target region of thalamocortical axons from the dorsal LGN give rise to an area of high bouton-density [8], enabling with high precision spatially accurate synaptic connections. Due to their particular morphology, resulting from relatively higher bifurcation ratios as compared to the long-range axonal projections which constitute the axonal trunk [75], these self-similar ramifications are called here *florets.* Figure 1 panel **A** visualizes the two different parts of the axonal arbor. Quantification of florets based on 3-D reconstructions of the axon happens in two steps: First, a mean-shift algorithm developed by Binzegger et al. [8] is used to find bouton-dense patches, which are candidate regions for florets. This process is repeated until all such clusters are identified. All axonal structures inside these clusters whose initial segment has come from *side-branching* are then defined as florets. A side-branch is defined as a branch that grows approximately perpendicular to the original direction of the axon, thereby distinguishing it from other bifurcations, which exhibit smaller branching angles between 20-80° [53, 75].

### 2.3 Quantitative global and individual analysis of florets

Various indicators have been proposed for quantification of neurite morphologies [7, 18, 67, 68]. We choose indicators based on properties of the entire dataset (global indicators) as well as indicators quantifying properties of individual florets (individual indicators). This allows us to account for overall governing growth rules as well as for individual floret morphology. The two criteria considered for optimization of the floret-generator are the global segment-length distribution calculated for all segments of the dataset, and a weighted asymmetry index calculated individually for each floret. Additionally we introduce eight further individual criteria which are not employed for optimization of the floret-generator to evaluate artificial florets. These criteria quantify mean and standard deviation of segment lengths, the number of segment lengths and the depth for each floret, as well as the non-weighted asymmetry value for each floret.

#### 2.3.1 Global segment-length distribution

Segment-lengths as well as all individual indicators discussed in this paper can be fully captured by 2-D representations of growth morphologies called *dendrograms*. As illustrated in Figure 2, dendrograms plot the segment-lengths along the x-axis while the branching topology is preserved. We distinguish here between different kinds of segments: the *root* segment, the *internal* segments, and *terminal* segments at the tips of florets. Together with its descendants a segment constitutes a *sub-floret.* The *depth* of the segment is the tree depth in the dendrogram: The root segment has depth one, while each bifurcation increments the depth of subsequent segments by one. As florets are binary trees, each bifurcation leads to exactly two new segments. These concepts are illustrated by dendrograms in Figure 2. For an estimation of the global segment-length distribution first a histogram of all 2060 segments from the dataset is constructed. Then a maximum-likelihood fit is used to calculate the best fitting distribution to the histogram. To determine the best fitting distribution, standard distributions are ranked by a Bayesian information criterion [37].

**Figure 2:**
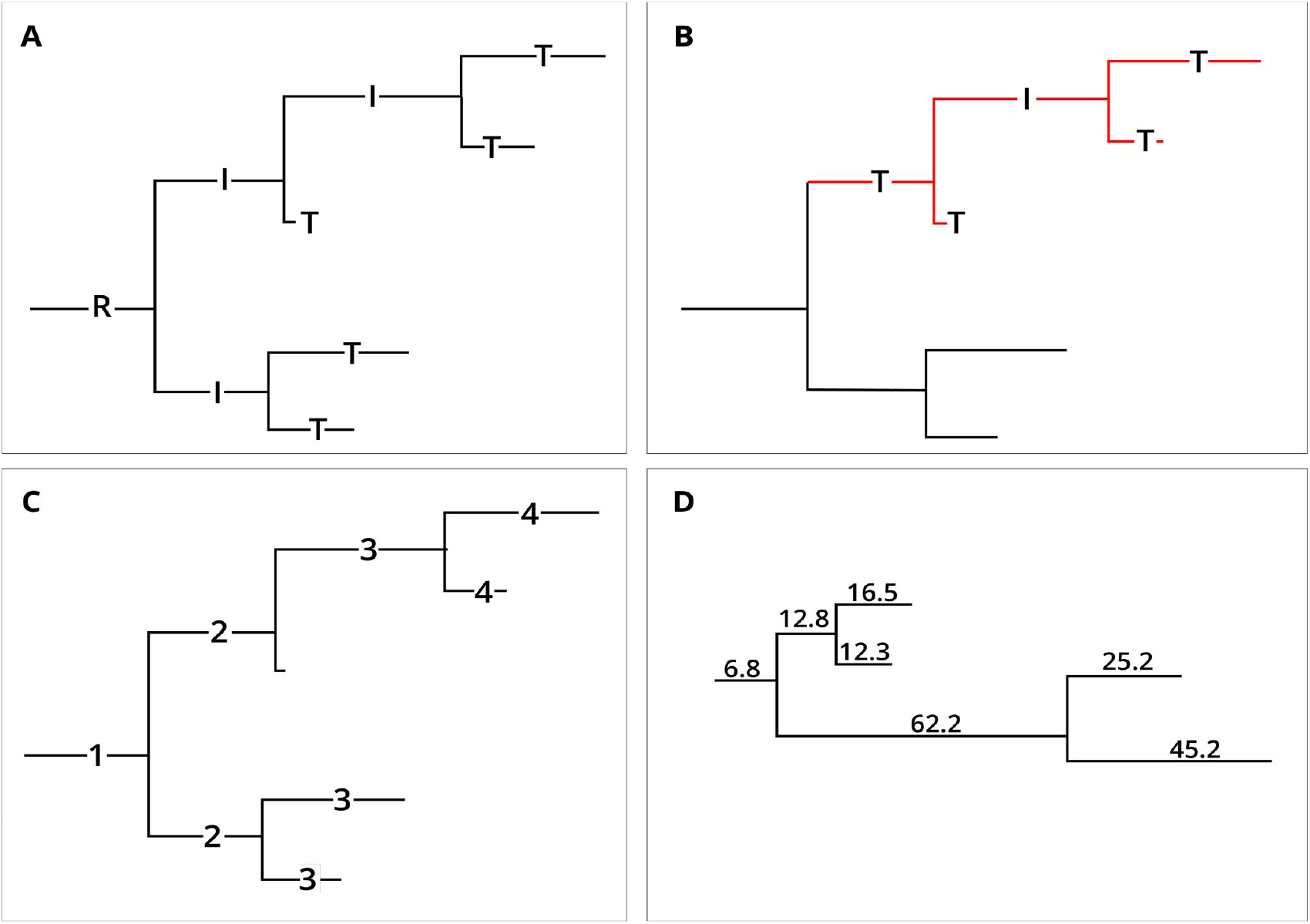
The indicators employed for evaluation of florets can be fully represented by dendrograms. (A) The dendrogram of a floret together with the different types of segments: Terminal segments (“T”), the root segment (“R”) and the internal segments (“I”). (B) An exemplary sub-floret is shown in red. (C) Each segment labeled by its depth. While the root segment has depth one, each subsequent bifurcation increments the depth by one unit. (D) A floret from the dataset which is perfectly symmetrical according to Colless’ asymmetry index (asymmetry index = 0), but has an asymmetry of 0.41 according to the weighted asymmetry index.

#### 2.3.2 Weighted asymmetry: A novel quantification for neurite morphology

We introduce here a novel asymmetry index to quantify the asymmetry of neurites, called the *weighted* asymmetry index. It combines topological and metrical criteria, with topology understood as the connectivity pattern of the segments [65]. This index is an extension of a classical asymmetry index for tree-structures originating in the work of Colless [11] and extensively used in studies on neurite branching patterns [67]. Colless’ index takes only the number of segments into account, independent from differences in segment lengths. Colless’ asymmetry index A as applied to binary neurites [67] can be defined as follows:

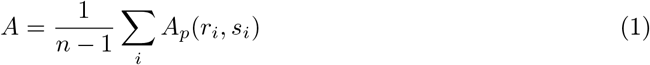

where *A_p_* denotes the asymmetry of each subtree, and *r_i_* and *s_i_* are the number of leaves in the left and right subtree originating from branching point *i. A_p_* is then computed as

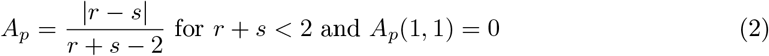

The weighted asymmetry index *A_w_* in contrast employs the length of segments to weigh the contribution of each subtree, while being equivalent to Colless’ index for binary trees with identical segment lengths. Let *F* be a floret with *n* segments *s_i_, i ∈ I* = {1,…,*n*}. Every segment *s_i_* defines exactly one sub-floret *F*(*s_i_*) containing *s_i_* as its root segment and all its descendants. Since every segment bifurcates into exactly two new segments, *F* has exactly 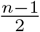 non-terminal sub-florets. For every such sub-floret we define *s_i_l__* and *s_i_r__* as the left and right sub-floret originating from *s_i_*. Furthermore, for every sub-floret *G* of *F* we define |*G*| as the number of segments in the sub-floret, *t*(*G*) as the number of terminal segments in *G*, and 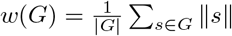 as the mean segment length in *G*. We can now define the weighted asymmetry index *A_pw_* for a sub-floret *F*(*s_i_*) as

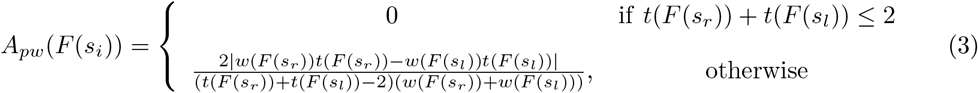

Here, we compare the number of segments in all left and right subflorets, but weigh this number by the length of the respective segments. Finally, this value is normalized to lie in the unit interval [0, 1], as is the case for Colless’ index. Similar to (1), the weighted asymmetry index of an entire floret *F* is computed as the mean of its non-trivial sub-florets’ asymmetry, where the sum goes over all sub-florets of *F*:

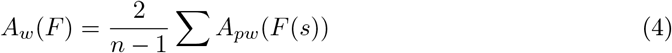

For both weighted and Colless’ asymmetry index a value of zero corresponds to a fully symmetrical and a value of 1 to a completely asymmetrical floret. Figure 2 shows a floret from the dataset, which is fully symmetrical according to Colless’ but not according to the weighted asymmetry index.

#### 2.3.3 Further individual quantification criteria for florets

In addition to weighted asymmetry index and segment-length distribution, a number of other indicators are employed for the evaluation of generated florets. These additional indicators all quantify individual floret morphology. To compare generated and real florets, a distribution of these indicators over the entire dataset is calculated. Importantly, these criteria are not used for optimization of the floret-generator. A good match between real and generated florets in terms of these criteria can hence serve as a further validation of the floret-generator. The eight additional indicators employed for evaluation are displayed and explained in Table 1.

**Table 1:**
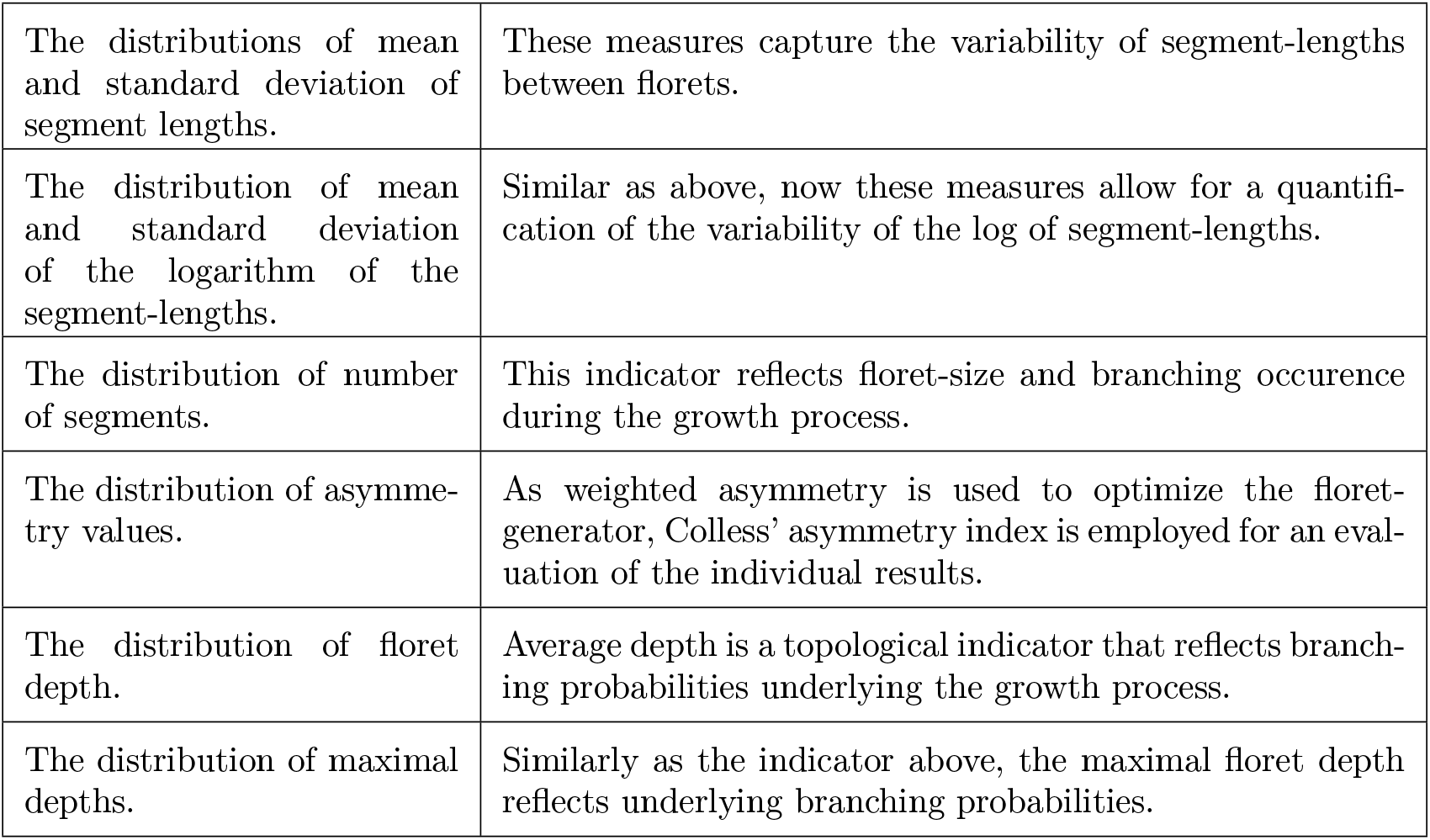
Individual criteria for evaluation of generated florets. These criteria are not employed for the optimization of the floret-generator. The criteria capture metric (segment-length) and topological (asymmetry, depth) properties, as well as floret size (number of segments).

#### 2.3.4 Optimization of the floret-generator model

Mathworks’ built-in genetic algorithm (GA) from the Global Optimization Toolbox, version 3.4.1., is used to optimize the parameters of the floret-generator [36]. This is achieved by minimizing the divergence between original and generated dataset. In particular, the divergence is calculated between normalized histograms of the biological and the generated data, whereby histograms are constructed with MATLAB’s built-in histogram method [39]. We seek to minimize the divergence of segment-length and weighted asymmetry distributions between real and generated data to match global as well as individual statistics of the florets. This gives rise to the following objective function the genetic algorithm employs:

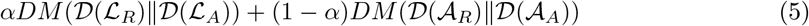

where 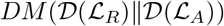 denotes the divergence measurement (*DM*) between the distribution of segment-lengths of the real dataset 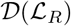 and of the generated dataset 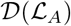 is the divergence between the distribution of weighted asymmetry indices of the real dataset 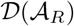 and of the generated dataset 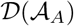. *α* is a scaling factor which is determined empirically. As divergence measure the symmetric Jensen-Shannon (JS) divergence is employed [34]. Its symmetry ensures that an optimal trade-off for fitting peak as well as tail of the distribution is found. In a first step the GA randomly generates initial solutions. In subsequent steps, ten percent of solutions with the best fitness values (“elites”) are guaranteed to be passed on to the next generation, while additional solutions for the next generation are created through crossover and mutation. The algorithm halts if a maximum number of generations, here set to ten times the number of parameters, is reached or if the weighted average change over 50 generations falls below 10^−6^.

## 3 Results

We present the global and individual statistics of the biological dataset, and compare these to the statistics of the generated florets. The floret-generator model is optimized with regards to the segment-length and weighted asymmetry distribution of the real data, however further statistical criteria are used to assess the generated florets. In addition, we assess if the MATLAB implementation leads to equivalent results as the Cx3D implementation. As the latter operates include biophysical interactions with the environment, a successful translation into this environment underlines the biological plausibility of our model. Lastly, using again global and individual statistics, we compare the floret-generator to the Galton-Watson model.

### 3.1 Global segment-length distribution of the floret dataset

The global segment-length distribution (Figure 3) of the biological data, containing a total of 2060 segments, displays a unimodal distribution with a visible lack of short segments. The best fit is indeed achieved by a log-normal distribution which is a longtailed, unimodal, right-skewed distribution. By definition, the logarithm of a log-normal distribution is a normal distribution. However, the segment-length distribution from the biological dataset displays a shift to the left. Indeed only the logarithm of the shifted segment-distribution passes the Kolmogorov-Smirnov (KS) test for normality at the 5% significance level, as without the shift, the logarithm of the segment-length distribution displays a skewness of −0.2726. A shift parameter of *γ* = −2.9157 and parameters *μ* = 3.52 and *σ* = 1.03 are obtained for the global segment-length distribution [38].

**Figure 3:**
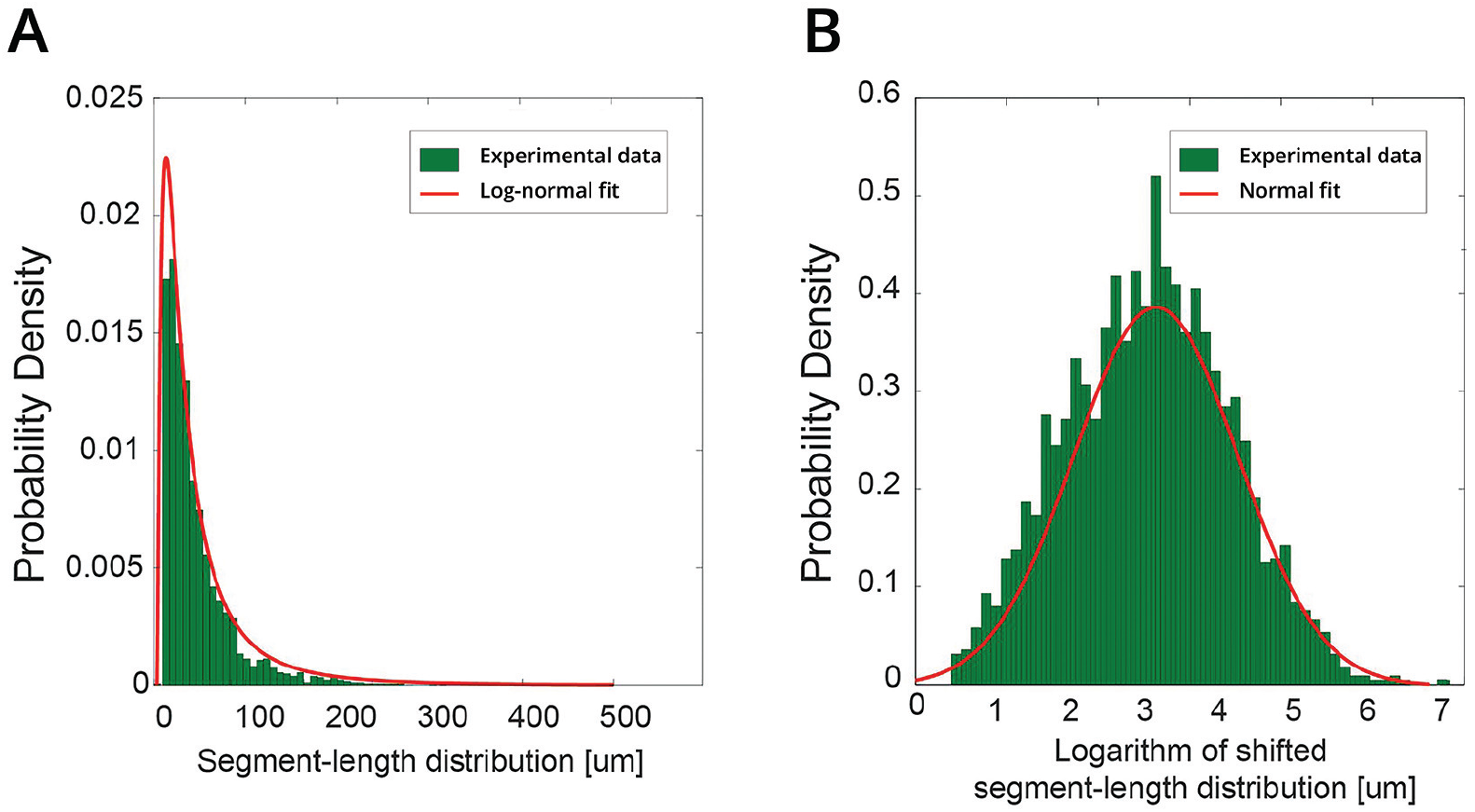
(A) Histogram of all 2060 segment-lengths of the dataset and best maximum-likelihood fit. The best fit is achieved by a log-normal distribution. (B) Histogram of the logarithm of the shifted segment-length distribution. The histogram displays now a low skewness of –0.0636 and passes the KS test for normality at the 5% significance level.

### 3.2 Individual statistics of the floret dataset

The individual statistics reveal that the florets of the dataset are highly variable in their morphology. Key statistical properties of the florets are presented in Table 5, while Figure 4 displays the eight individual criteria as defined in Table 1 and not employed for optimization of the floret-generator. 260 out of the 426 florets of the dataset are trivial florets. These florets have only one segment, and accordingly are fully symmetrical and have a depth of 1. In contrast, non-trivial florets of the dataset have on average 10.84 branches and an asymmetry of 0.39, with an outlier floret consisting of 232 branches. While the mean segment-length is around 50 microns for both trivial and non-trivial florets, the dataset contains florets with a mean segment-length of up to 440 microns, and the range of segment-lengths goes from 1 micron up to almost 910 microns. The statistics further reveal that not only the morphological variability between florets is high, but that highly variable metrics can be found within individual florets as well. This is captured by the distribution of standard deviations of segment-lengths and log segment-lengths, which display that a considerable fraction of florets contain segments of highly variable lengths.

**Figure 4:**
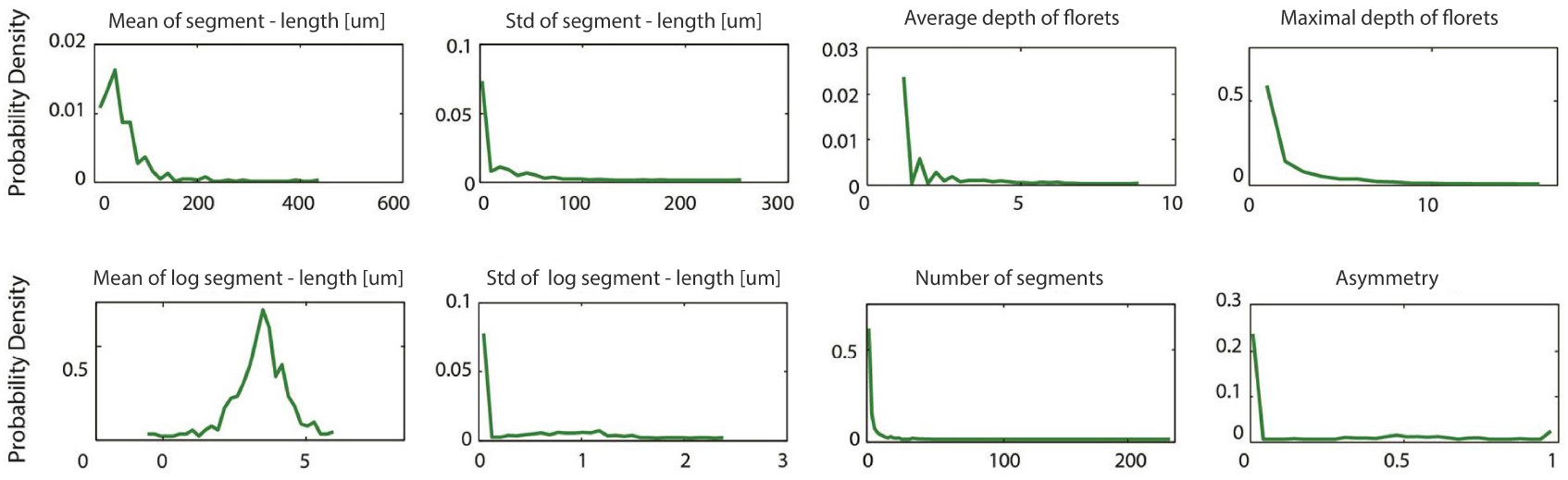
Quantification of individual statistics of the florets. A large fraction of florets consist of only one segment. This is reflected in the plots with asymmetry and depth distributions having high densities around 0 and 1 respectively. Non-trivial florets however contain a higher number of segments, and show a large variability in their segment-lengths, as illustrated by the distribution of the standard-deviations of segment-lengths and log segment-lengths. The morphological variability of the florets is also visible in the distribution of asymmetries, which covers the entire range of the index.

### 3.3 The floret-generator model

We introduce here the algorithm of the floret-generator in pseudo-code in Table 2, its parameters are presented in Table 3. This algorithm is implemented in MATLAB, which allows for the optimization of the parameters with a genetic algorithm. The optimized model is later ported into Cx3D, where the morphological asymmetry is implemented as resulting from the interaction of the growth process with chemotaxic cues in its environment.

**Table 2:**
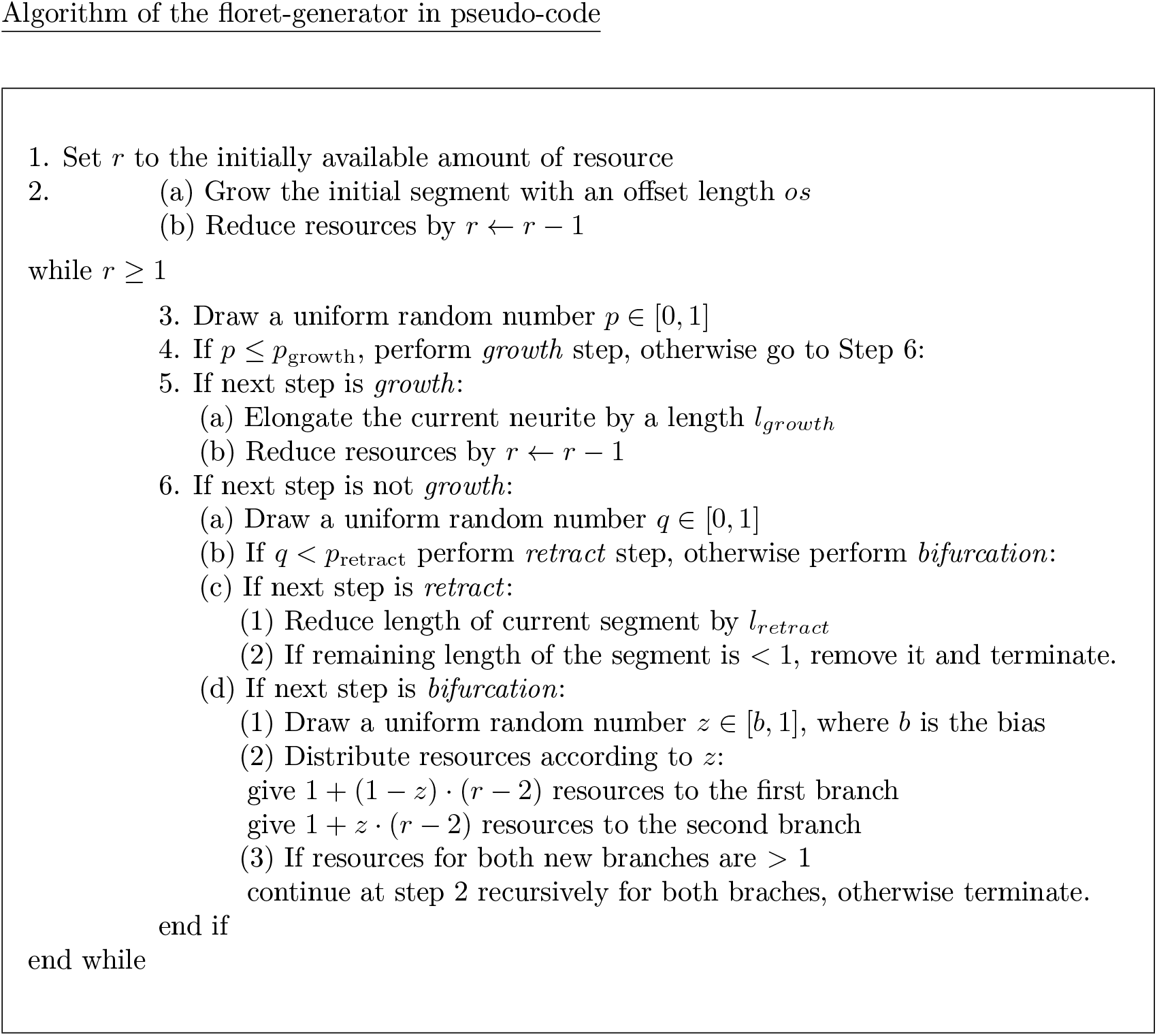
The algorithm of the floret-generator in pseudo-code. As an initial step, an offset-length (os) is grown. Subsequently, further growth, branching or retraction can occur. Resource is reduced by one unit after each growth step, and a bias parameter (b) decides on the distribution of resource after each bifurcation. Growth and retraction length as well as resource are sampled from a gamma distribution. The parameters of the gamma distribution as well as all other parameters of the model are optimized by a genetic algorithm.

The floret-generator we propose here is an extension of the Galton-Watson branching process [29]. The Galton-Watson branching process can be described as follows: At each step, three possible actions are available – growth, bifurcation and halting. Each growth step leads to the addition of a constant length, which does not change during the entire growth process. Similarly, the probability of occurrence for any of the three actions is constant during the entire growth process.

The proposed floret-generator model allows for growth, bifurcation and additionally for retraction, and makes use of an abstract resource parameter to control the halting of the algorithm. In particular, each floret is assigned an initial resource budget *r*. The resource budget is sampled for each floret from a gamma distribution whose parameters have been optimized with the genetic algorithm based on the entire data set. Each growth step requires one unit of resource, which is subtracted from the initial budget after the addition of a new segment. When the growth cone bifurcates, the resource is randomly distributed between the left and right child branches. The distribution happens according to a random sample from [*b*, 1], where *b* denotes the *bias* parameter. As the bias parameter is optimized in the interval [0.5,1], the resource distribution can be asymmetrical.

In line with the resource parameter, the growth and retraction lengths *l_growth_* and *l_retract_* are uniformly sampled from a gamma distribution. Again, the parameters of this distribution are optimized by a genetic algorithm based on the entire data set. The model employs a growth *offset*: each floret grows an initial minimum length *os*. All subsequent growth processes follow the same rules - at every step each growth cone may either grow, branch, or retract with the corresponding probability. If the retracted length is longer than the grown length, the respective branch is removed. The algorithm terminates when the resource falls below two units. Importantly, the proposed model stochastically creates artificial florets without requiring global coordination or communication, in accordance with biological principles of self-construction [4, 5, 27, 75].

#### 3.3.1 Parameters and optimization of the floret-generator

The model parameters are presented in Table 3. The parameters of the floret-generator are optimized with a genetic algorithm with regards to equation 5. While the parameters are optimized by taking the entire dataset into account, during the growth process each floret randomly samples from these optimized distributions.

**Table 3:** Parameters of the floret-generator. With exception of growth and retraction probabilities, which are sampled from the unit interval, the parameters are optimized by a genetic algorithm in their individual ranges, within which the genetic algorithm seeks an optimum over several iterations of its search process. The objective function of the genetic algorithm seeks to minimize the divergence of segment-length and weighted asymmetry distributions between real and generated data. As divergence measure the symmetric Jensen-Shannon divergence is employed.

|  |  |
| --- | --- |
| Growth and retraction parameters $[rs_{Sh}, rs_{Sc}]$ | Elongation and retraction lengths $[l_{growth}, l_{retract}]$ are drawn from a gamma distribution with shape and scale parameters both optimized in the range of $[0,100]$. |
| Resource parameter $[r]$ | For every floret, the amount of available resource is sampled from a gamma distribution. The gamma parameters for resource are both optimized in the range of $[0,20]$. |
| Growth and retraction probabilities $[p_{growth}, p_{retract}]$ | The probability of growth and retraction are both sampled uniformly from $[0,1]$. |
| The bias parameter $[b]$ | The bias parameter is optimized in the range of $[0.5,1]$. It lower bounds the amount of resource one descendant receives after bifurcation. |
| The offset parameter $[os]$ | The offset parameter decides the initial length which is grown before all other steps. It is optimized in the range of $[1,2]$ |

#### 3.3.2 Segment-length distribution of the generated floret dataset

The best fit for the global segment-length distribution is presented in Figure 5, panel A. The generated segment-length distribution is based on 500 generated florets. Displayed is the generated distribution with the lowest Jensen-Shannon divergence of 0.011. It is clearly visible that the segment-length distribution matches the peak as well as the long tail of the segment-length distribution of the real dataset. The generated distribution is also capable of accounting for the short segments found in the original dataset. In summary, the generated segment-lengths cover the entire range of the original segment-lengths but also reflect the variability of these, hence reproducing the shifted log-normal distribution found in the original dataset. This is confirmed by a Kolmogorov-Smirnov test for the equality of the distributions at the 5% significance level. Table 4 contains the parameters of the floret-generator as optimized by the GA. The optimized bias parameter is close to 0.5. This indicates a rather equal distribution of resource after branching events, and the generation of on average rather symmetrical florets. This observation matches the above observed mean asymmetry of 0.15 for the entire dataset. The scale parameter of retraction is comparatively high, although retraction probability is lower than growth probability. Hence while retraction appears less frequently, retraction lengths are on average larger than grown segment-lengths. Scale and shape parameters of the gamma distribution for resource are rather large, close to the upper bound of their optimization interval of [0, 20].

**Figure 5:**
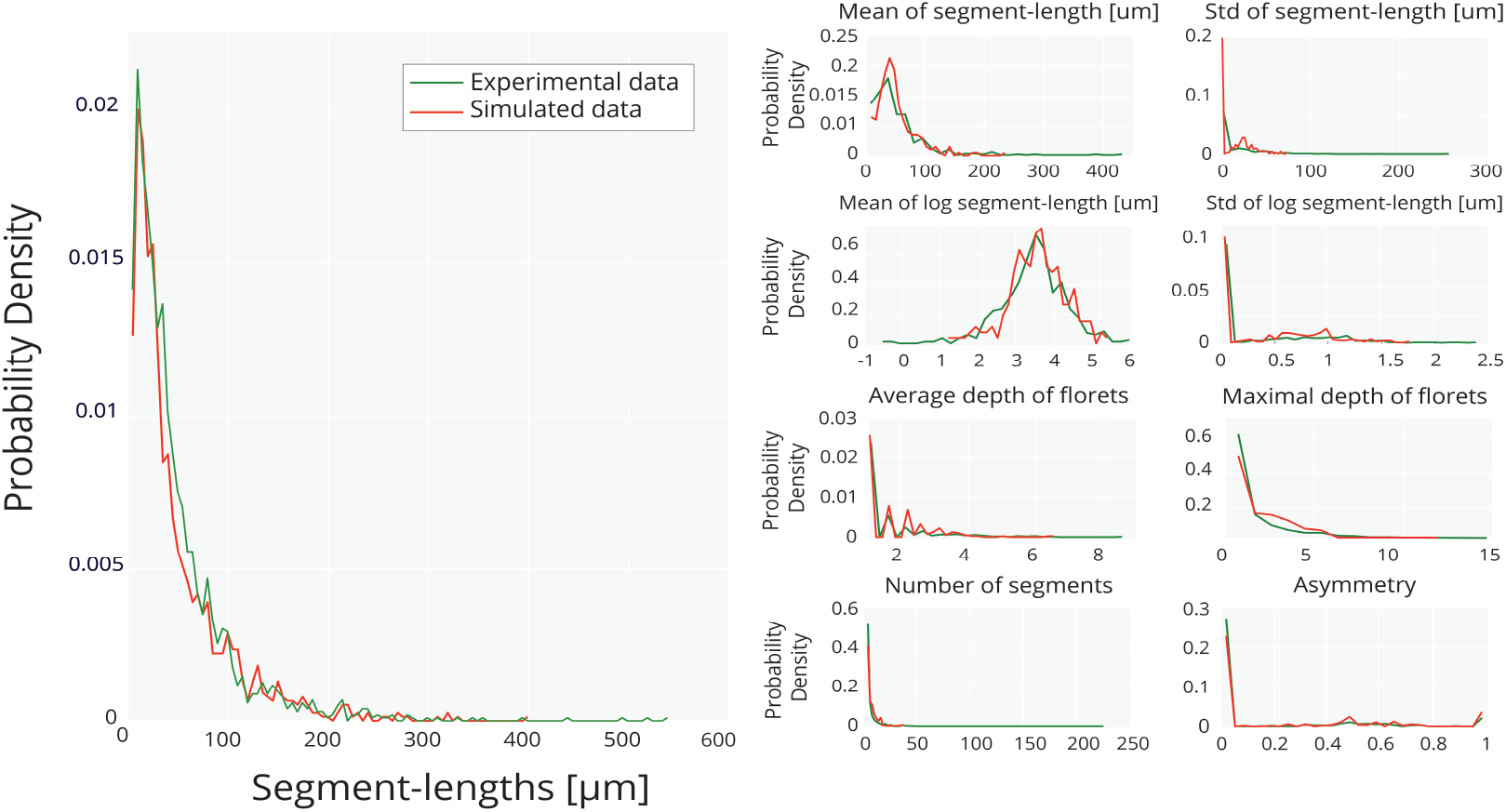
(A) The best generated and the real global segment-length distributions in comparison. The low Jensen-Shannon divergence of 0.011 indicates a good overall fit. As can be seen, the generated segment-length distribution matches peak as well as tail of the real segment-length distribution, and is capable of generating the log-normal form. The equality of the distributions is confirmed by a Kolmogorov-Smirnov test at the 5% significance level. (B) Comparison of the generated and real individual statistics. The generated statistics capture the fraction of trivial florets which are completely symmetric and have depth one. Concurrently, the generated florets are also reproducing the entire ranges of the indicators.

**Table 4:** Parameters for the offset-variation with retraction and a gamma distribution for growth and retraction lengths: The gamma distribution with shape and scale parameter for growth lengths (g_*Sh*_, g_*Sc*_), retraction lengths (r_*Sh*_, r_*Sc*_) and resource (rs_*Sh*_, rs_*Sc*_). The growth probability (p_*g*_). The retraction probability (p_*r*_). The bias parameter (b). The offset set parameter (os). The bias parameter is close to 0.5, indicating a rather equal distribution of resource after branching events. The scale parameter of retraction is comparatively high. Although retraction occurrence is lower than growth occurrence, retracted lengths are on average larger than growth lengths.

| $g_{Sh}$ | $g_{Sc}$ | $r_{Sh}$ | $r_{Sc}$ | $rs_{Sh}$ | $rs_{Sc}$ | $p_g$ | $p_r$ | $b$ | $os$ |
| --- | --- | --- | --- | --- | --- | --- | --- | --- | --- |
| 2.24 | 5.52 | 2.28 | 53.80 | 19.66 | 18.10 | 0.67 | 0.46 | 0.54 | 1.26 |

#### 3.3.3 Individual statistics of the generated floret dataset

A comparison of the individual statistics of generated and real florets from the best generated run is shown in Figure 5, panel B. As can be seen the generated florets capture the statistical properties of trivial florets found in the dataset. This becomes apparent upon inspection of depth, segment number and asymmetry distributions, where the generated population captures the large fraction of symmetrical florets with only one segment and a depth of 1. The same distributions reveal that the generated florets concurrently capture a wide range of these attributes. The good fit of individual statistics is also reflected by the fact that the generator matches the distributions of the standard deviation of segment-lengths and log segment-lengths. This demonstrates that the floret-generator can create highly variable florets as well as florets which display highly variable segments, as found in the real dataset. With the exception of depth and standard deviation of segment length distributions, all individual statistics pass the Kolmogorov-Smirnov test for equality at the 5% significance level.

### 3.4 The floret-generator in Cx3D

Using the model parameters as optimized with MATLAB, we next translate the floret-generator into Cx3D (“Cortex 3D”), a simulation environment for neural development [73]. Cortical development in Cx3D is based on the idea of the development of the brain through locally autonomous processes, in accordance with principles of cortical self-construction. A successful translation accordingly underlines the agreement of the proposed model with these principles. With regards to our growth model, the autonomous processes in pysical 3D space can be interpreted as the growth cones steering neurite outgrowth. Assuming that growth cones follow chemotaxic cues [19, 22, 41], we can model the asymmetry observed in the florets now by introducing such a substance into the neurites’ extracellular matrix. This is done by modelling the concentration of the substance as constant along the x-and y-axis, but linearly changing along the z-axis, the florets’ direction of growth. This constant slope is equal to the bias parameter introduced in Table 2, whereby again 1 > *bias* > 0.5.

Subsequently the gradient vector *υ* of the substance equals (0, 0, *bias*), when *z* is in the aforementioned interval. The norm of this vector *υ* equals the bias parameters as introduced earlier, i.e. ||*υ*|| = *bias*. Hence, resource allocation is now governed by the norm of the gradient of the substance: When bifurcating, the neurite senses and “extracts” the gradient of the surrounding substance. We assume that noise is being sampled uniformly from the interval [(1 – ||*υ*||), 1]. This can be interpreted as noise intrinsical to gradient detection through the growth cones [43]. Adding noise to the norm of the gradient vector we match exactly our usage of the bias parameter for the allocation of resources as introduced in Table 2. In order to achieve growth into the direction of the gradient we allocate the larger fraction of resource to the segment enclosing a smaller angle with the gradient. In this manner, the asymmetrical growth of neurites is modelled through an interaction of the neurite with its extracellular environment. The parameters of the CX3D-floret-generator correspond to the floret-generator parameters as described above in Table 3.

#### 3.4.1 Simulation of the floret-generator in Cx3D

We demonstrate empirically that the floret-generator can be ported into Cx3D by comparing the segment-length distributions and individual statistics generated by the algorithm described in Table 2 and implemented in MATLAB to those generated in the Cx3D environment as described in Section 3.4. For this comparison we generate 500 florets in each of the implementations, using the previously optimized parameters of the floret-generator presented in Table 3. In the MATLAB implementation the distribution of resource after bifurcations is decided by randomly sampling from the interval [b,1], with the bias parameter *b* being optimized by the GA. In contrast in Cx3D we model resource distribution as a function of extracellular gradients. As visible in Figures 7 the segment-length distributions and the individual statistics of the two implementations show a nearly perfect fit, demonstrating a successful translation of the floret-generator into the Cx3D simulation environment. All the statistics of the MATLAB and the Cx3D pass the Kolmogorov-Smirnov test for equality at the 5% significance level. The slight divergence between the two models can be attributed to the stochasticity of the modelled growth-processes.

**Figure 6:**
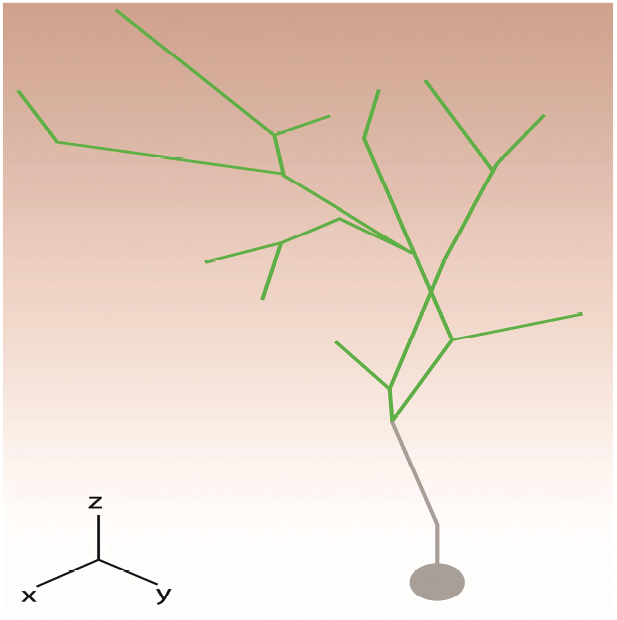
A floret generated in Cx3D. An extracellular gradient is shown in the background (red). The gradient changes linearly along the z-axis. After bifurcation, the norm of the gradient plus an additional noise component determine how resource is distributed to the two offspring growth cones.

**Figure 7:**
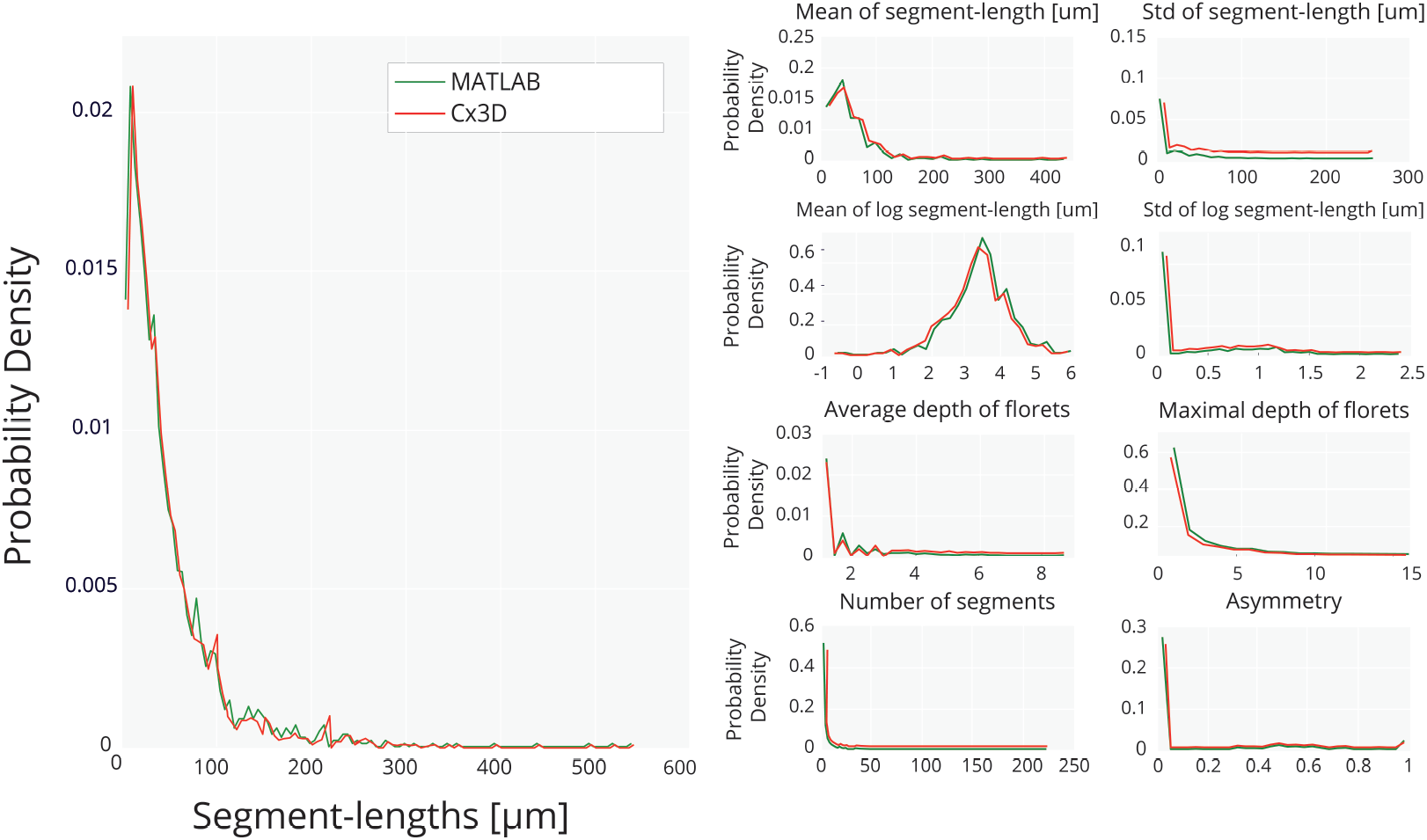
(A) Comparison of the global segment-length distribution from the floret-generator implemented in MATLAB and in the physical simulation environment provided by Cx3D. The distributions are based on 500 generated florets in each implementation. Although the distribution of resource is modelled in Cx3D as an interactive process influenced by extracellular gradients, the global statistics generated by both implementations proves to be equivalent. (B) Comparison of the individual statistics for the floret-generator implemented in MATLAB and in Cx3D. The individual statistics are calculated based on 500 generated florets in each implementation. The individual statistics show a close match, and both global and individual statistics pass the Kolmogorov-Smirnov test for equality at the 5% significance level. The slight divergence in the results can be attributed to the stochasticity of the floret-generator.

### 3.5 The floret-generator in comparison to the Galton-Watson model

The Galton-Watson model is the oldest and most commonly employed branching process. Here we fit a Galton-Watson model to the biological dataset, and compare the global and individual statistics from this model to the results from the floret-generator and the biological dataset. During the growth process of the Galton-Watson model, three possible actions are available at each step: Bifurcation, growth or halting. The respective probabilities p_*b*_, p_*gr*_, p_*st*_ remain constant during the entire growth process and fully determine the segment-length distribution [7, 29]. The length added at each growth step is constant as well, and taken together these properties lead to an exponential segment-length distribution. In accordance with the work of Binzegger et al. [7] which is partially based on the same dataset, we chose a segment-length of 1. As the probabilities sum up to 1, we optimize only p_b_, p_gr_ of the Galton-Watson model with respect to the objective function defined in equation 5. We employ the same genetic algorithm as previously discussed, and create as before 500 artificial florets. In line with Binzegger et al. [7], we use the following condition to prevent infinite growth [29]:

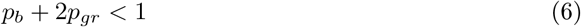

The optimized Galton-Watson model has a growth probability p_*gr*_ of 0.98 and a branching probability of p_*b*_ of 0.0031, comparable to the results of Binzegger et al. [7]. Figure 8 displays the global and individual results of an optimized Galton-Watson process. As shown in previous work [7], this model fails to capture the log-normal distribution of the segment-lengths. In particular, it is not capable to recreate the lack of small segment-lengths as observed for the biological data. Its exponential segment-length distribution leads to a Jensen-Shannon divergence of 0.026. This discrepancy between artificially generated florets from the Galton-Watson model and the biological data is also apparent in the individual statistics, displayed in Table 5. As opposed to the floret-generator, the Galton-Watson model fails at recreating branches with accurate lengths. This can be again explained by the failure of the Galton-Watson model to recreate the lack of shorter branches of the biological dataset, which can not be compensated by the short tail of its generated segment-length distribution, and the lack of the possibility for retraction. With the exception of the asymmetry distribution, indeed the comparison of Galton-Watson generated and biological distributions fail the Kolmogorov-Smirnov test for equality at the 5% significance level, for global and individual statistics.

**Figure 8:**
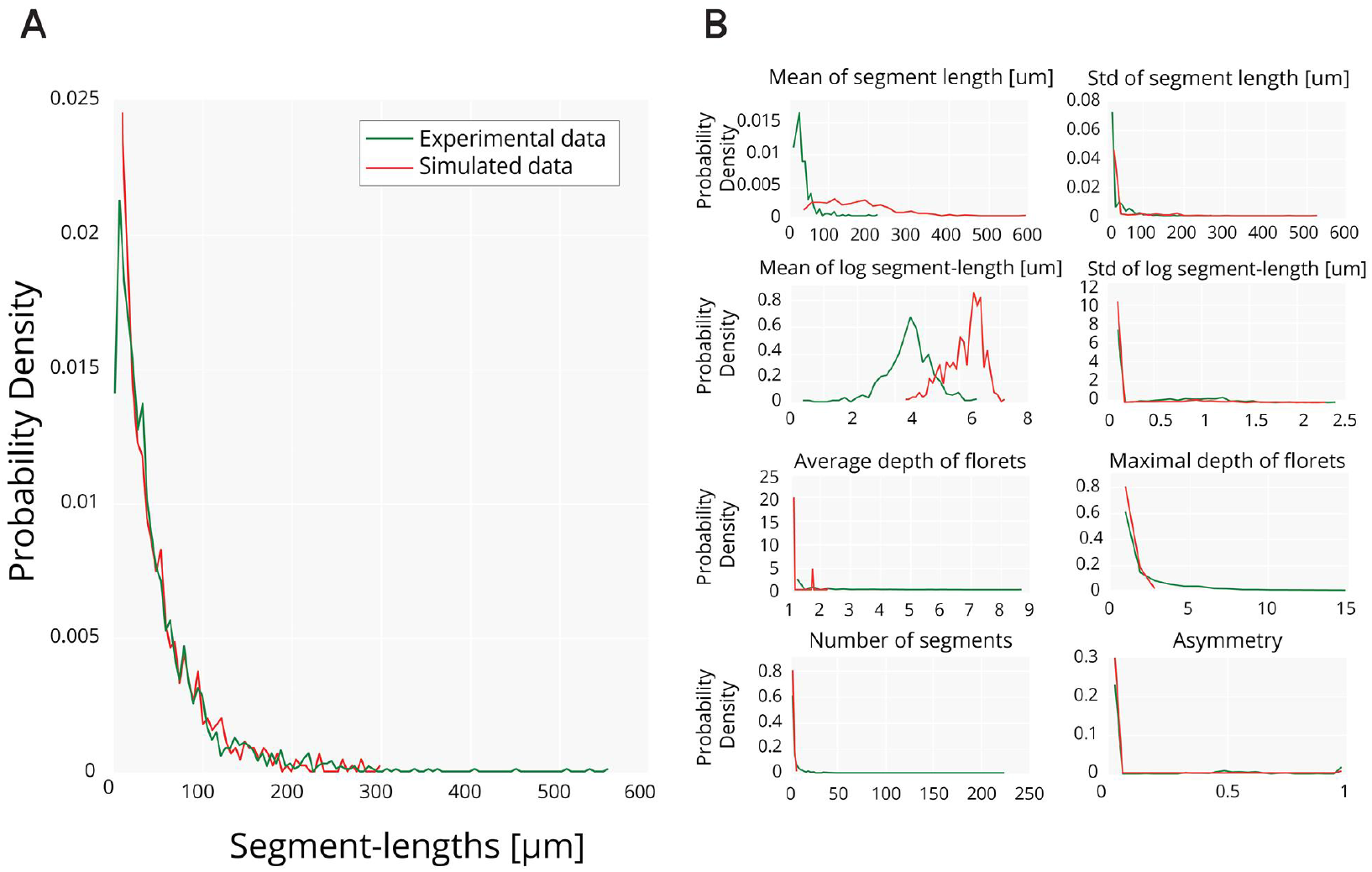
(A) The best generated from the Galton-Watson model, and the real global segment-length distributions in comparison. The Jensen-Shannon divergence between the two distributions is with 0.026 higher than for the previously presented floret generator. This can be explained by the model’s failure to recreate the lack of short segments, and the tail of the biological data. The distributions fail the Kolmogorov-Smirnov test for equality at the 5% significance level. (B) Comparison of the generated and real individual statistics. The lack of the short segment can not be recreated by the Galton-Watson distribution. Accordingly, the generated segment-lengths do not match the biological data as adequately as the floret-generator is capable of. Indeed with the exception of the asymmetry distribution, all individual distributions fail the Kolmogorov-Smirnov test for equality at the 5% significance level.

**Table 5:** Statistical properties of the floret dataset from cat thalamic axons and the three generated datasets. Where not indicated otherwise, the criterion is calculated over the entire dataset. The floret-generator, both in MATLAB and Cx3D, matches the biological data best. The Galton-Watson model diverges markedly from the statistics of the dataset with regards to the generated segment-lengths. This can be ascribed to its failure of reproducing peak and tail of the segment-length distribution of the biological data. The average asymmetry of the biological data equals 0.15, a rather low value which can be explained by the large number of trivial florets present in the dataset. The average asymmetry of the non-trivial florets equals 0.39. This measure can be matched well by both floret-generator and Galton-Watson model. Avg.: Average; SD: Standard deviation; NT: Non-trivial floret; T: Trivial floret.

|  | Real data | MATLAB | Cx3D | Galton-Watson |
| --- | --- | --- | --- | --- |
| Avg. segment-lengths | 49.9 $\mu\text{m}$ | 52.57 $\mu\text{m}$ | 53.1 $\mu\text{m}$ | 312.5 $\mu\text{m}$ |
| Avg. segment-lengths NT | 48.89 $\mu\text{m}$ | 45.49 $\mu\text{m}$ | 45.8 $\mu\text{m}$ | 134.58 $\mu\text{m}$ |
| Avg. segment-lengths T | 50.55 $\mu\text{m}$ | 68.2 $\mu\text{m}$ | 64.1 $\mu\text{m}$ | 355.55 $\mu\text{m}$ |
| Avg. SD segment-lengths | 15.99 $\mu\text{m}$ | 23.3 $\mu\text{m}$ | 23.3 $\mu\text{m}$ | 19.06 $\mu\text{m}$ |
| Avg. SD segment-lengths NT | 41.04 $\mu\text{m}$ | 36.1 $\mu\text{m}$ | 36.2 $\mu\text{m}$ | 97.84 $\mu\text{m}$ |
| Avg. depth | 1.69 | 1.77 | 1.78 | 1.13 |
| Avg. depth NT | 2.79 | 2.2 | 2.2 | 1.67 |
| Avg. asymmetry | 0.15 | 0.24 | 0.21 | 0.01 |
| Avg. asymmetry NT | 0.39 | 0.35 | 0.35 | 0.02 |

## 4 Discussion and conclusions

“Patchy” regions of thalamic arborizations have been observed in various species, and these dense local ramifications are thought to preserve the spatial resolution of the associated receptive fields [2], and to enable dense synaptic connections for the ocular dominance columns [32, 33]. Our results demonstrate that the proposed generative model recreates the individual morphologies and global statistics of such ramifications of cat thalamic afferents in visual area 17 with remarkable accuracy. Importantly, the model achieves these results while being based on locally autonomous growth processes, in accordance with developmental principles of the brain [5, 48, 75] and in the tradition of mechanistic modelling approaches in line with these principles [2, 5, 7, 42].

By taking asymmetry and segment-length distributions into account, the floret-generator model however expands on previous mechanistic approaches to neurite modelling [57], and, in addition, surpasses the statistical accuracy of the Galton-Watson model, the best known and most employed branching model. For instance, the model is capable of matching the global shifted log-normal distribution of the segment-lengths almost perfectly. This is a stark improvement to the exponential segment-length distribution generated by the Galton-Watson model for the same dataset [7]. The log-normal distribution of the axonal segment-lengths stems from a distinct lack of short segments, which as Binzegger et al. [7] point out, is not an artefact of reconstruction, since such short segment-lengths could be easily seen in the light microscope. The difference in generated global statistics by the two models is crucial, since a good match to the statistics of experimentally observed data serves as a benchmark for the accuracy of inferred growth rules, and hence provides the basis for studies of the functionality of associated neuronal circuits. The good fit to the shifted log-normal segment-length distribution can be attributed to the imposed minimal length by an offset parameter, and the variable probabilities of the possible actions, sampled at the beginning of each growth process from a gamma distribution.

By capturing the metrical properties of the dataset, the global segment-length distribution can reveal characteristics about growth, branching and retraction probabilities as well as resource parameters which all impact metric properties of generated segments. Our models predicts for instance that retraction probabilities are lower than growth probabilities, while retracted lengths were larger than grown lengths. As neurite outgrowth is assumed to depend on tubulin, transported to the growing tip of the neurite [21, 47], conditioning it on the availability of a resource parameter further underlines the biologically plausibility of the proposed model. This modelling approach is also in contrast with neurite modelling approaches which are based on formal optimization considerations, and hence do not allow for insights into such mechanistic attributes of the growth process [9, 12, 13].

The good fit to the global statistics can be attributed to several further properties of the floret-generator. Together with an observed high expected value for resource, the inclusion of retraction contributes to the capability of the floret-generator to create highly diverse florets. A high resource budget can for instance create florets with numerous segments, while infrequent occurrences of retraction can prune some of these to smaller lengths. This leads to artificial florets with metrically highly variable segments, as present in the original dataset. In addition, the large expected value of resource together with a low branching probability -which implies that numerous steps of growth can occur before the next branching point - can be attributed for matching the tail of the global segment-length distribution. Retraction can in addition lead to the recreation of the smaller segment-lengths. Taken together this explains the capability of the floret-generator to match the peaks as well as the tails of the log-normal distribution.

Besides global statistics, individual neurite morphology is an important measure for identification and differentiation of neurites and neurons and can reveal the influence of external factors shaping the neurite [7, 21, 71]. Previous work has shown that topological asymmetry is a very discriminative criterion for branching patterns, while also reflecting the developmental history of these [67]. Accordingly a novel *weighted* asymmetry index was introduced here to allow for a finer control of individual statistics than the previously used quantification from Colless [7, 11, 71]. The individual statistics of artificial florets confirmed indeed an accurate recreation of the morphology of real florets. The generated individual statistics capture properties of the large fraction of trivial florets with only one segment and quantified as entirely symmetric found in the original dataset. Concurrently, the floret-generator accounts also for the more variable fraction of rather asymmetrical florets with a larger number of segments. The bias parameter of the model, optimized in the interval [0.5,1], proved to be rather low with a value of 0.54. This reflects the overall low asymmetry of 0.15 found in the original dataset, which can be attributed to the large fraction of trivial florets. It is important to note that the Galton-Watson model fails to model the fraction of small segments, and and so the individual statistics generated by this model are less in accordance with the experimental data.

With the goal to provide a biophysical interpretation for statistical observations, the floret-generator was ported into the simulation environment of Cx3D. As neurogenesis in this simulation environment operates on the basis of autonomous local processes, the successful translation of the floret-generator into Cx3D underlines its biological plausibility. In this simulation environment the bias parameter is interpreted as the gradient of an extracellular cue, linearly changing along the axis of growth. As such, the asymmetry of the florets is modelled as the result of the growth cones interaction with the environment. After bifurcation, the larger amount of resource was allocated to the branch enclosing a smaller angle with the extracellular gradient, reflecting the fundamental role of chemotaxic cues for axonal path finding [19, 22, 41]. Again, this implementation did not assume any kind of global coordination between the growth cones as opposed to previous work in the literature [69, 71]. Future work could expand the model by the inclusion of further biophysical constraints. Branching could for instance be explicitly modelled as depending on a resource parameter as it has been suggested that its occurrence is determined by the availability of resources such as tubulin and other proteins such as MAP2 [47]. It must be further noted that we had restricted ourselves to a two dimensional analysis of the florets. The inclusion of angles in Cx3D might lead to further conclusions about the underlying growth principles. Theoretical studies for instance suggest that wiring processes optimize branching angles with regards to a minimization of branching lengths [30, 51]. Furthermore, while the electrical activity of neurons is the topic of intense studies, it is rather rarely studied in the context of neurite development. However, neurite growth, branching and retraction during development are strongly influenced by electrical activity [54, 63, 72], and in turn determine the signaling properties of neuronal circuits [40, 52]. A further extension of our work could accordingly be the inclusion of electrical signaling activity, in line with the work of Bauer et al. [5] where this activity emerges based on developmental self-construction of the circuitry.

Our model is a clear improvement to previous models for local axonal growth, which were based on the Galton-Watson branching process [7, 45]. The proposed model can recreate floretal statistics not only with higher accuracy, it is also a biologically plausible model and as such provides a solid basis for future analysis of the functionality of associated neuronal circuits. Importantly, the high statistical accuracy of our model is achieved while only making use of a small set of simple rules. This demonstrates how axonal morphologies of high variability can be the product of a developmental process governed by the repeated instantiation of the same basic, genetically encoded rules. Formally, the presented floret-generator model falls into the category of *Bellman-Harris* processes, a generalization of the Galton-Watson process [31]. Analysis of Bellman-Harris processes with respect to neurite outgrowth might allow for further valuable deductions regarding neurite growth processes. In addition, future studies should test the model on other experimentally collected datasets, and integrate the floret-generator in broader simulations of cortical development [4, 5, 75].

## Acknowledgements

The authors thank Rodney J. Douglas, Kevan A.C. Martin, Andreas Hauri, John C. Anderson, Peter Jagers, Tom Binzegger and Nuno DaCosta for valuable advice throughout the work. Conflict of Interest: None declared.

